# The comparison of single-cell RNA sequencing platforms based droplets

**DOI:** 10.1101/2024.06.16.599202

**Authors:** Linyan Wang, Yulong Zhong

**Affiliations:** Eye Center, The Second Affiliated Hospital, School of Medicine, Zhejiang University, Zhejiang Provincial Key Laboratory of Ophthalmology, Zhejiang Provincial Clinical Research Center for Eye Diseases, Zhejiang Provincial Engineering Institute on Eye Diseases, Hangzhou, Zhejiang, China; Guangdong Provincial Key Laboratory of Bone and Joint Degenerative Diseases, Department of Physiology, School of Basic Medical Sciences, Southern Medical University, Guangzhou 510515, China

## Abstract

Single-cell sequencing enables to reveal cellular heterogeneity and discover new cellular subpopulations. In terms of the strategy of single-cell sequencing, the main methods are based with combinatorial index, microwell and microfluidic. Due to the simplicity, methods based droplets are widely used for single-cell sequencing for multi-omics. Therefore, in order to facilitate researchers to choose a suitable platform to meet their application scenarios, we compared several commercial platforms: the Chromium X platform of 10x Genomics, the MobiNova-100 platform of MobiDrop, the SeekOne platform of SeekGene, and the C4 platform of BGI. Based the comprehensive assessment of the data analysis, the Chromium X platform shows a excellent performance, closely followed by MobiNova-100 platform.

**One-Sentence Summary:** As droplet-based single-cell sequencing platforms, Chromium X and MobiNova-100 have comparable data performance.

## Introduction

Cells are the units of organisms and there is significant heterogeneity among cells(*1*). When analyzing large tissue samples consisting of millions of cells, it is difficult to distinguish heterogeneity between individual cells and identify cells that may play an important role in development and disease progression (*2-5*). In order to enable the studies of cellular heterogeneity, single-cell sequencing technology was developed with 96-well plates in 2009 (*6*), but this strategy is escalated due to be laborious and complicated (*7-9*). In 2015, droplet-based single-cell sequencing was developed, which easily captures cells with high throughput and the process is simple and convenient (*10-11*). As many advantages appear to droplet-based single-cell RNA-seq sequencing technology, there are many excellent solutions successfully commercialized, such as the Chromium X series of 10x Genomics, the MobiNova-100 platform of MobiDrop, the SeekOne platform of SeekGene, and the C4 platform of BGI (*12-15*). Since many solutions are available for single-cell sequencing, it is especially important for researchers to find a suitable solution (*16-18*).

In order to illustrate the performance of different platforms, we performed experiments and compared the results of captured cell number, captured gene number, cellular clustering, cell type annotation, differential gene expression and cell signal pathway on the four platforms above. The results show that the comprehensive performance Mobinova-100 platform is close to the performance of Chromium X series. In terms of the significance of differential genes, the performance of MobiNova-100 is particularly excellent.

## Result

### The quality control of four platforms based droplets

The identification of distinct cell types through the clustering of scRNA-seq profiles stands out as a key application. The datasets from Peripheral Blood Mononuclear Cells (PBMCs) encompasses a variety of cell types, providing an opportunity to evaluate and contrast different methodologies for this particular scenario. We compared several commercial platforms with PBMCs.

Sensitivity is an important metric to evaluate scRNA-seq methods. It represents the ability of the technology to capture RNA molecules with a small amount of input. We used the UMI per cell and the number of genes captured per cell to evaluate the capture efficiency of the method. Overall, the 10x Genomics platform captured the most UMIs and number of genes per cell (median genes=2085, median UMIs=5532) of the four platforms. Among them, MobiDrop and SeekGene had close UMIs median and gene median with 10x Genomics platform (respectively 4365,1747 for MobiDrop; respectively 5172,1782 for Seekgene). The BGI platform had the fewest genes and UMIs per cell (Table.1).

**Table 1.** The comparison of data quality control.

| Platform | 10x Genomics | MobiDrop | SeekGene | BGI |
| --- | --- | --- | --- | --- |
| Mapping rate | 92.2% | 94.71% | 96.11% | 94.11% |
| Cell number | 15417 | 17670 | 4949 | 5837 |
| Median gene number | 2085 | 1747 | 1782 | 661 |
| Median UMI | 5532.5 | 4365 | 5172 | 1049 |
| Exon ratio | 53.7% | 59.15% | 52.37% | 53.7% |
| Intron ratio | 30.2% | 28.05% | 31.30% | 28.1% |
| Total gene ratio | 83.9% | 87.2% | 83.67% | 81.8% |

Note: The median gene number and median UMI are filtered from four single-cell sequencing platforms with the same parameter.

As we analysis to integrate different data sets, we can comparison between different methods for estimation of doublets rate. We found the highest doublets rate for 10x Genomics, slightly lower for Mobidrop, and comparable and low doublets rate for SeekGene and BGI. The higher doublets rate of 10x Genomics and Mobidrop can be attributed to the larger number of cells (*19*).

Note: The median gene number and median UMl are filtered from four single-cell sequencing platforms with the same parameter.

### Clustering and annotation of total PBMCs

Sctransform normalization was used to normalize the data and Harmony was used to integrate all the collected data (*20-21*). PCA (Principal Component Analysis) and UMAP (Uniform Manifold Approximation and Projection) were performed to reduce dimension for the data visualization. The cell number of SeekGene and BGI platform is extremely low than 10x Genomics and MobiDrop platforms, especially not enough cells that falls into some specific clusters. Finally, cells were annotated into 6 cell types: B cells, CD4^+^ T cells, CD8^+^ T cells, Monocytes, NK cells and DCs, consistent with previous studies (Fig.1) (*22*).

**Fig. 1.**
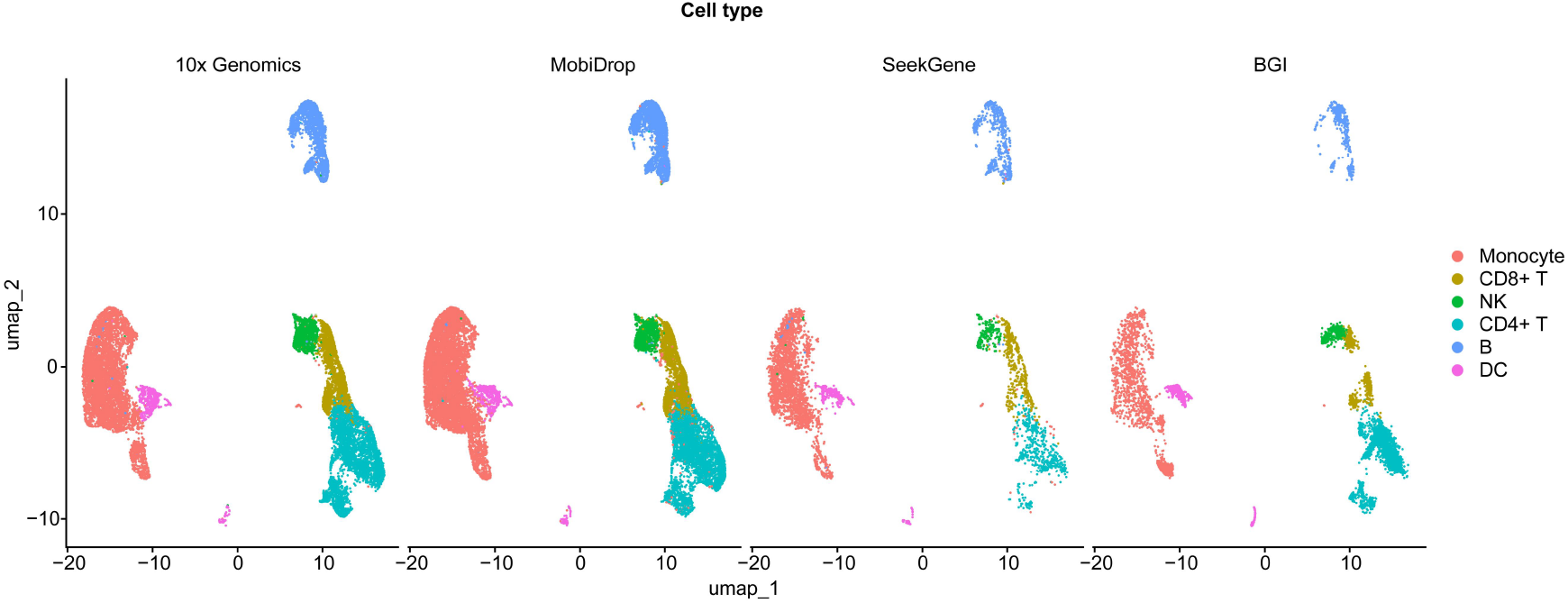
clustering and annotation of captured cells with different platforms. Single-cell RNA-seq was performed with human PBMC cells through four platforms. Clustering and cell annotation was carried out to obtain 6 cell populations in total: B cells, CD4 T cells, CD8 T cells, monocytes, DC cells, NK cells.

Based the analysis, it was observed that every cell type was identified with each respective dataset. However, methods varied in the ability to distinguish cell types, in the proportion of cell types recovered. As expected, methods were more difficult to distinguish transcriptionally related cell types, such as CD4 T cells, CD8 T cells and natural killer cells. From the UMAP plot, we observed that for identifying CD8 T cells, BGI performed slightly better, while the other three methods performed equally well. We also found that for most cell types, although MobiDrop, 10x Genomics and SeekGene were not the best, their performance was relatively consistent and robust. BGI is less effective at identifying individual cell types, such as monocytes. Since all libraries for each experiment were generated from the same sample, we evaluated the consistency of the different methods in terms of the proportion of cells assigned to each cell type in a single experiment. In summary, all methods recovered all cell types, but the proportion of different cell types varied.

The data showed that MobiDrop had the most similar cell ratios to the 10x Genomics platform. The differences in the cell ratios of CD4 T cells and monocytes captured by the SeekGene platform were poor compared to the other platforms and the differences in the B cells captured by the BGI platform were poor compared to the other platforms (Fig. 2A). After de-batch effect, those results were comparable to analyze the performance of clustering by average silhouette width (ASW) (Fig. 2B).

**Fig. 2.**
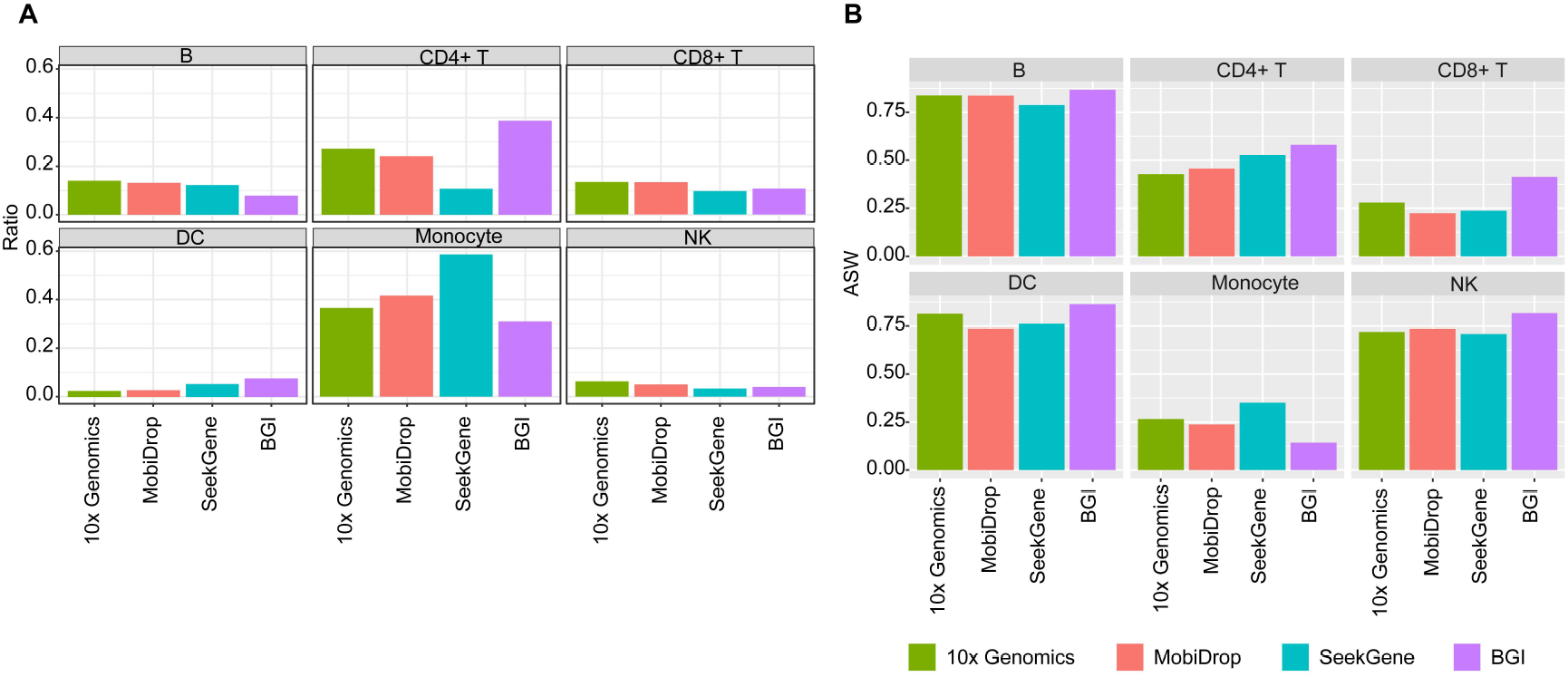
Proportion of captured cells with different platforms. (A) Statistics on the percentage of cells captured using different platforms. (B) Comparison of ASW values for different cell populations.

### Comparison of marker gene’s expression and high variation genes

To emphasize the accuracy of the clustering, we analyzed the expression of marker genes of the captured cell by each platform. The results showed that the strong signals of cell marker genes appear to MobiDrop and SeekGene platforms, while the 10x platform was relatively weak to detect marker gene expression and the BGI platform captured no obvious signal of marker genes. Among them, the MobiDrop platform performed well in B cells and DC cells with the high percentage and strong expression of marker genes. However, when using the 10x platform, the captured signal of T-cell maker genes appeared to be weak (Fig. 3A).

**Fig. 3.**
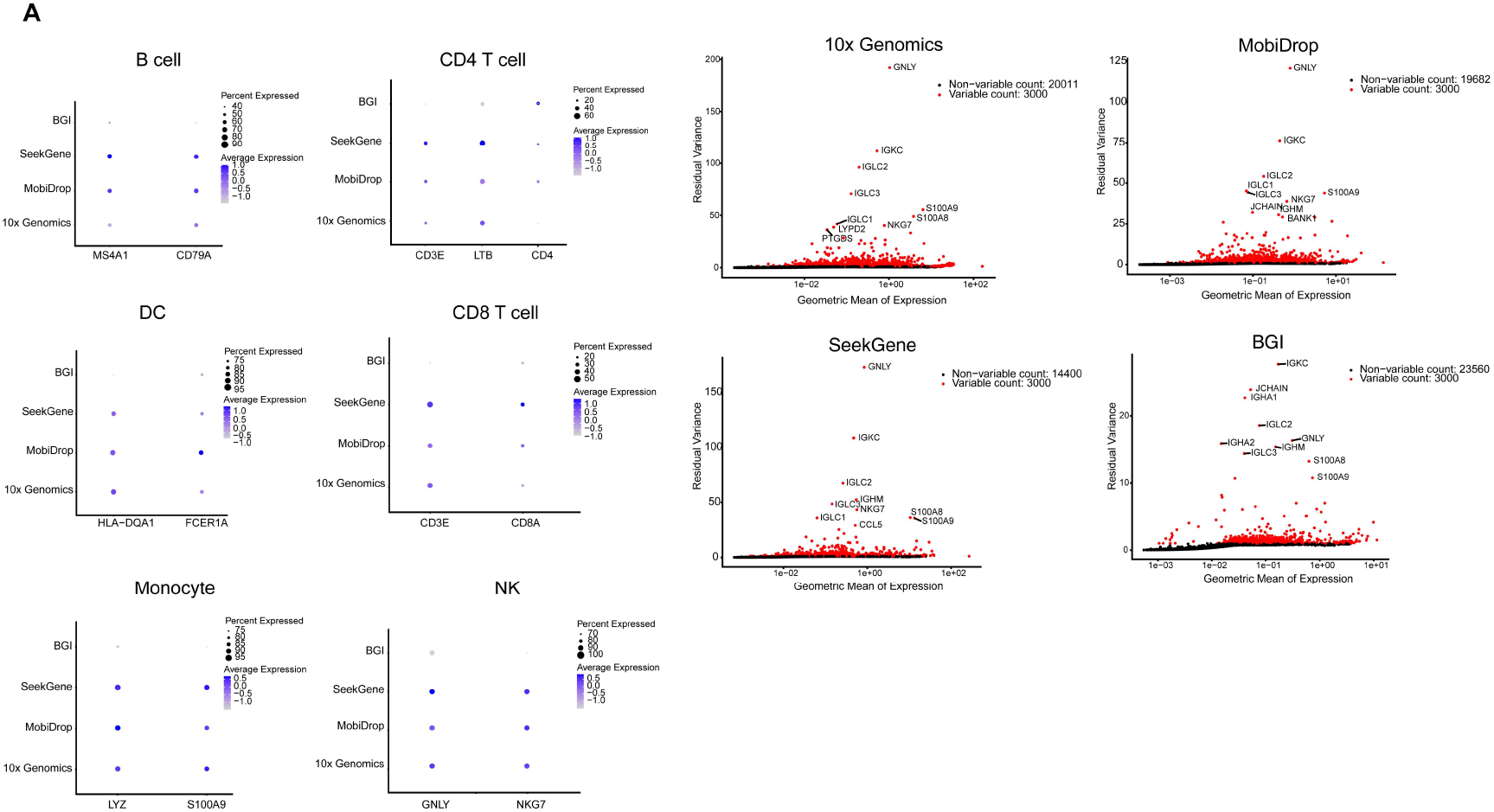
The analysis of high varaition genes. (A) Proportion and significance of marker genes captured with different platforms. (B) Expression of high variation genes with different platforms.

At the same time, we did a count of barcode for the four platforms used in this study. Based on the comparison of the results, we can find that the top 10 highly variable genes of MobiDrop, 10X, and SeekGene are mostly overlapping, in the order of GNLY, IGKC, IGLC2, IGLC1, IGLC3, S100A9, and NKG7, whereas the top 10 highly variable genes of BGI are partially overlapping, in the order of IGKC, IGLC2, GNLY, IGLC3, S100A9 (Fig. 3B).

The data above suggests that the performance of MobiDrop, 10x Genomics and SeekGene platforms are comparable with the estimation of marker gene expression and high variation genes.

### The capture of signal pathways in PBMCs

The importance of signaling pathways in peripheral blood mononuclear cells (PBMCs) lies in their critical role in regulating immune responses. PBMCs, which include lymphocytes (T cells, B cells, and NK cells) and monocytes, are key components of the immune system. The signaling pathways within these cells control their activation, differentiation, and function in response to various stimuli, such as infections, inflammation, and autoimmune reactions. Understanding these pathways is essential for elucidating the mechanisms of immune responses and disease pathogenesis.

We did enrichment analyses of the signaling pathways for each cell type in response to the data quality of the different platforms (Fig.4). The results showed that the MobiDrop, 10x Genomics and SeekGene platforms captured stronger signals, and the BGI platform was the least effective. Meanwhile, for CD4^+^ T cells, the performance of SeekGene platform was not stable enough.

**Fig. 4.**
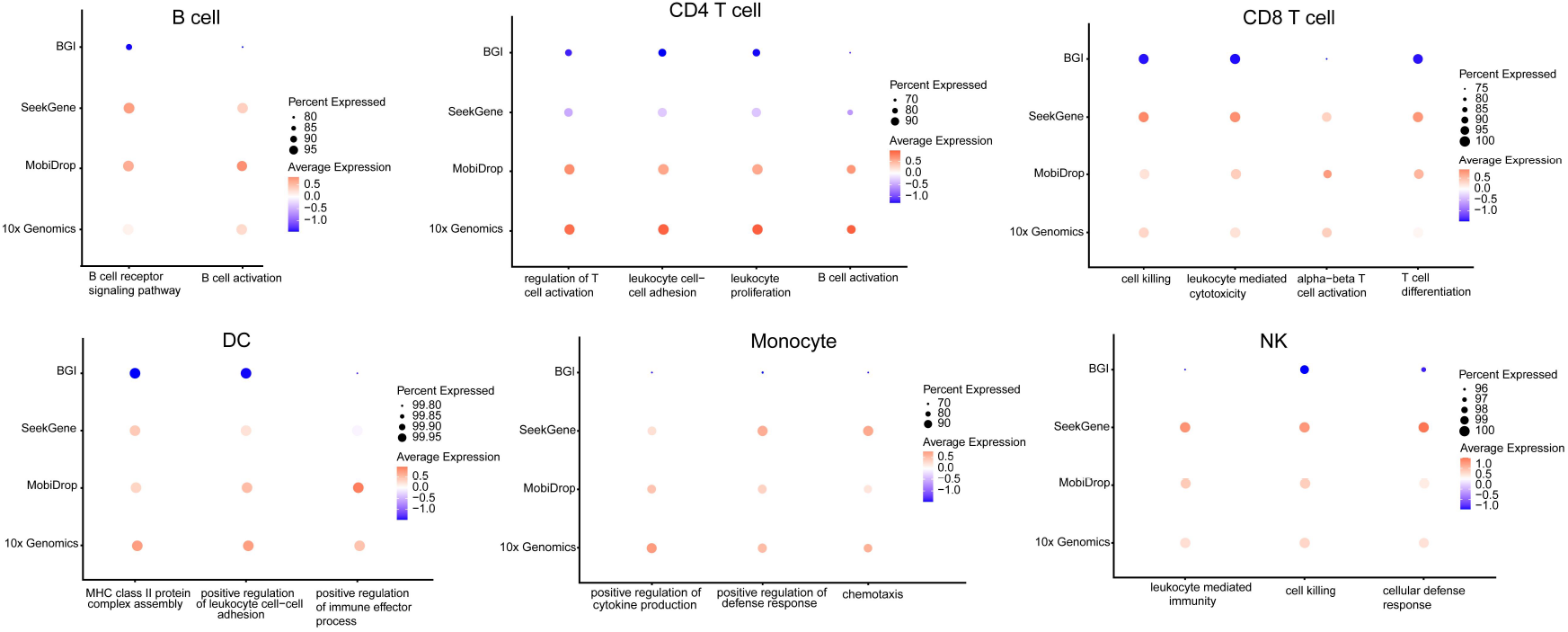
Captured signal pathways according to each cell type. Enrichment analysis of signalling pathways specific to each cell subpopulation

## Discussion

There are also low-throughput scRNA-seq methods that sort cells or use a mouth pipette to pick off cells into a single well of a 96-well plate. However, these methods can only analyze tens to hundreds of single-cell transcriptomes at a time. In the field of single-cell transcriptome sequencing, there are more microfluidic-based methods that simplify the operation and show great advantages.

In this study, we systematically compare four high-throughput single-cell sequencing platforms via droplet-based methods, including the Chromium X platform of 10x Genomics, the MobiNova-100 platform of MobiDrop, the SeekOne platform of SeekGene, and the C4 platform of BGI. As far as captured gene number is concerned, the 10x Genomics platform has the best performance, while MobiDrop has the best performance in terms of captured cell number and gene mapping rate. The performance of 10x Genomics and MobiDrop is comparable in terms of the effect of cell clustering and cell type ratio. In terms of the significance of marker genes and differential genes, MobiDrop and SeekGene show the best performance. As droplet-based single-cell sequencing platforms, Chromium X and MobiNova-100 reveal comparable data quality.

By comparing the different platforms in this study, we are able to choose the method that is suitable to our research scenario. For the detection of a rare subpopulation, choose a low-throughput method to load pre-purified cells. We can also load more cells in a high-throughput platform to obtain more information for all loaded cell subpopulations. Of course, cost should also be taken into account. When the number of cells is less than 100, a low-throughput method may be a better choice, otherwise, a high-throughput platform will perform better. In addition, the platform should be versatile that can be used to integrate with multi-omics to analyze gene expression and gene regulation through different dimensions.

## Acknowledgments

We thank Linyan Wang for critical comments on this manuscript.

## Author contributions

Y.Z. conceived and designed the study. Y.Z. designed and performed all experiments. Y.Z. performed the computational analyses. Y.Z. provided technical support and wrote the paper with L.W.. All authors participated in data discussion and interpretation.

## References and Notes

[1] Gordon S, Taylor PR. Monocyte and macrophage heterogeneity. Nat Rev Immunol. 2005;5(12):953–964.

[2] Dagogo-Jack I, Shaw AT. Tumour heterogeneity and resistance to cancer therapies. Nat Rev Clin Oncol. 2018;15(2):81–94.

[3] Papalexi E, Satija R. Single-cell RNA sequencing to explore immune cell heterogeneity. Nat Rev Immunol. 2018;18(1):35–45.

[4] Hou Y, Guo H, Cao C, et al. Single-cell triple omics sequencing reveals genetic, epigenetic, and transcriptomic heterogeneity in hepatocellular carcinomas. Cell Res. 2016;26(3):304–319.

[5] Zeng Q, Mousa M, Nadukkandy AS, et al. Understanding tumour endothelial cell heterogeneity and function from single-cell omics. Nat Rev Cancer. 2023;23(8):544–564.

[6] Tang F, Barbacioru C, Wang Y, et al. mRNA-Seq whole-transcriptome analysis of a single cell. Nat Methods. 2009;6(5):377–382.

[7] Ramsköld D, Luo S, Wang YC, et al. Full-length mRNA-Seq from single-cell levels of RNA and individual circulating tumor cells [published correction appears in Nat Biotechnol. 2020 Mar;38(3):374]. Nat Biotechnol. 2012;30(8):777–782.

[8] Picelli S, Björklund ÅK, Faridani OR, Sagasser S, Winberg G, Sandberg R. Smart-seq2 for sensitive full-length transcriptome profiling in single cells. Nat Methods. 2013;10(11):1096–1098.

[9] Hashimshony T, Senderovich N, Avital G, et al. CEL-Seq2: sensitive highly-multiplexed single-cell RNA-Seq. Genome Biol. 2016;17:77. Published 2016 Apr 28.

[10] Macosko EZ, Basu A, Satija R, et al. Highly Parallel Genome-wide Expression Profiling of Individual Cells Using Nanoliter Droplets. Cell. 2015;161(5):1202–1214.

[11] Klein AM, Mazutis L, Akartuna I, et al. Droplet barcoding for single-cell transcriptomics applied to embryonic stem cells. Cell. 2015;161(5):1187–1201.

[12] Kalucka J, de Rooij LPMH, Goveia J, et al. Single-Cell Transcriptome Atlas of Murine Endothelial Cells. Cell. 2020;180(4):764-779.e20.

[13] Dai H, Zhu C, Huai Q, et al. Chimeric antigen receptor-modified macrophages ameliorate liver fibrosis in preclinical models. J Hepatol. 2024;80(6):913–927.

[14] Cai G, Hua Z, Zhang L, et al. Single-cell transcriptome analysis reveals tumoral microenvironment heterogenicity and hypervascularization in human carotid body tumor. J Cell Physiol. 2024;239(4):e31175.

[15] Chen C, Guo Q, Liu Y, et al. Single-cell and spatial transcriptomics reveal POSTN+ cancer-associated fibroblasts correlated with immune suppression and tumour progression in non-small cell lung cancer. Clin Transl Med. 2023;13(12):e1515.

[16] Wang Q, Xiong H, Ai S, et al. CoBATCH for High-Throughput Single-Cell Epigenomic Profiling. Mol Cell. 2019;76(1):206-216.e7.

[17] Xiong H, Luo Y, Wang Q, Yu X, He A. Single-cell joint detection of chromatin occupancy and transcriptome enables higher-dimensional epigenomic reconstructions. Nat Methods. 2021;18(6):652–660.

[18] Han X, Wang R, Zhou Y, et al. Mapping the Mouse Cell Atlas by Microwell-Seq [published correction appears in Cell. 2018 May 17;173(5):1307]. Cell. 2018;172(5):1091-1107.e17.

[19] Zheng, G. X. Y. et al. Massively parallel digital transcriptional profiling of single cells. Nat. Commun. 8, 14049 (2017).

[20] Hao, Y. et al. Dictionary learning for integrative, multimodal and scalable single-cell analysis. Nat. Biotechnol. 42, 293–304 (2024).

[21] Korsunsky, I. et al. Fast, sensitive and accurate integration of single-cell data with Harmony. Nat. Methods 16, 1289–1296 (2019).

[22] Ding, J. et al. Systematic comparison of single-cell and single-nucleus RNA-sequencing methods. Nat. Biotechnol. 38, 737–746 (2020).

[23] McGinnis CS. et al. DoubletFinder: Doublet Detection in Single-Cell RNA Sequencing Data Using Artificial Nearest Neighbors. Cell Syst. 2019;8(4):329-337.e4.

