## Supplementary Materials for "The comparison of single-cell RNA sequencing platforms based droplets"

**Materials and Methods**

**Cell preparation of PBMCs**

Frozen healthy human PBMCs were purchased from ORIBIOTECH. Frozen primary cells were thawed according to the manufacturer's instructions.

**Single-cell RNA-seq library preparation and sequencing (MobiDrop, MobiNova-100)**

The PBMCs were loaded into microfluidic chip of Chip A Single Cell Kit v2.1 (MobiDrop (Zhejiang) Co., Ltd., cat. no. S050100201) to generate droplets with MobiNova-100(MobiDrop (Zhejiang) Co., Ltd., cat. no.A1A40001). Each cell was involved into a droplet which contained a gel bead linked with up to millions oligos (cell unique barcode). After encapsulation, droplets suffer light cut by MobiNovaSP-100(MobiDrop (Zhejiang) Co., Ltd., cat. no.A2A40001) while oligos diffuse into reaction mix. The mRNAs were captured by cell barcodes with cDNA amplification in droplets. Following reverse transcription, cDNAs with barcodes were amplified, and a library was constructed using the High Throughput Single Cell 3'RNA-Seq Kit v2.0 (MobiDrop (Zhejiang) Co., Ltd., cat. no. S050200201) and the 3' Single Index Kit(MobiDrop (Zhejiang) Co., Ltd., cat. no. S050300201) . The resulting libraries were sequenced on an Illumina NovaSeq 6000 System.

**Single-cell RNA-seq library preparation and sequencing (10x Genomics, Chromium X)**

Single-cell 3'-RNA-Seq samples were prepared using single cell V3.1 reagent kit and loaded in the Chromium Controller according to standard manufacturer protocol (10x Genomics, PN-120237) to capture cells. Briefly, PBMCs were encapsulated in nanodroplets (GEMs) using a microfluidic device. These GEMs were generated combining barcoded single cell 3' V3.1 gel beads, a master mix that contains the reverse transcription (RT) reagents, the single cells, and partitioning oil onto the Chromium Next GEM Chip. After cell lysis, RNAs were captured on the gel beads coated with oligos containing an oligo-dT, unique molecular identifiers (UMIs) and a specific barcode. Incubation of the GEMs leads to the production of barcoded full-length cDNA from poly-A mRNA. After reverse transcription, GEMs were broken and cDNAs were purified with silane magnetic beads. Then, barcoded full-length cDNA was amplified by PCR to generate enough material for library construction. Amplified cDNA was purified again, and cDNA quality control was assessed by capillary electrophoresis (Bioanalyzer, Agilent) before the preparation of the libraries. Finally, libraries were prepared using a fixed proportion of the total cDNA. Enzymatic fragmentation and size selection were used to optimize the cDNA amplicon size. During the GEM incubation, the read 1 primer sequence was added to the molecules. At this step, P5, P7, a sample index, and the read 2 primer sequence were added via end repair, A-tailing, adaptor ligation and PCR. The final libraries that contain the P5 and P7 primers used in Illumina bridge amplification were sequenced on an Illumina NovaSeq 6000 System.

**Single-cell RNA-seq library preparation and sequencing (BGI, C4)**

Single-cell RNA-seq libraries were prepared using C4 scRNA-seq kit (BGI). Barcoded mRNA capture beads, droplet generation oil and the single-cell suspension were loaded into the corresponding reservoirs on chip for droplet generation. The droplets were gently removed to the collection vial and placed at room temperature for 20 minutes. Droplets were then broken and collected by the bead filter (BGI). The supernatant was removed, and the bead pellet was resuspended with 100 μl RT mix. The mixture was then thermal cycled as follows: 42 °C for 90 minutes, 10 cycles of 50 °C for 2 minutes, 42 °C for 2 minutes. The bead pellet was then resuspended in 200 μl of exonuclease mix and incubated at 37 °C for 45 minutes. Afterward the PCR master mix was added to the beads pellet and thermal cycled as follows: 95 °C for 3 minutes, 13 cycles (for nuclei 19 cycles) of 98 °C for 20 s, 58 °C for 20 s, 72 °C for 3 minutes, and finally 72 °C for 5 minutes. Amplified cDNA was purified using 60 μl of AMPure XP beads. The cDNA was subsequently fragmented to 400-600bp with NEBNext dsDNA Fragmentase (New England Biolabs) according to the manufacturer's protocol. Indexed sequencing libraries were constructed using the reagents in the C4 scRNA-seq kit following the steps: (1) post fragmentation size selection with AMPure XP beads; (2) end repair and A-tailing; (3) adapter ligation; (4) post ligation purification with AMPure XP beads; (5) sample index PCR and size selection with AMPure XP beads. The barcode sequencing libraries were quantified by Qubit (Invitrogen). All libraries were further prepared based on BGISEQ-500 sequencing platform. The DNA nanoballs (DNBs) were loaded into the patterned nanoarrays and sequenced on the BGISEQ-500 sequencer using the following read length: 41 bp for read 1, 100 bp for read 2, and 10 bp for sample index.

**Single-cell RNA-seq library preparation and sequencing (SeekGene, SeekOne)**

The single-cell RNA-seq libraries were prepared using SeekOne Digital Droplet Single Cell 3′ library preparation kit (SeekGene). An appropriate number of PBMCs were combined with the reverse transcription reagent and subsequently introduced into the sample well within the SeekOne DD Chip S3 (Chip S3). Following this, barcoded hydrogel beads (BHBs) and partitioning oil were separately dispensed into their corresponding wells within Chip S3. Once emulsion droplets were generated, reverse transcription was executed at a temperature of 42 ℃ for a duration of 90 min, followed by inactivation at 85 ℃ for 5 min. Subsequently, cDNA was extracted from the broken droplets and subjected to amplification through PCR reactions. The resulting amplified cDNA product underwent a series of steps including purification, fragmentation, end repair, A-tailing, and ligation to sequencing adaptors. Finally, indexed PCR was performed to amplify the DNA, which represented the 3' polyA region of expressing genes and also included the cell barcode and unique molecular index. The indexed sequencing libraries were subjected to purification using SPRI beads, followed by quantification using quantitative PCR. The libraries were then sequenced on an Illumina NovaSeq 6000 platform with PE150 read length.

**Single-cell RNA-seq data processing**

We implemented several quality control measures to eliminate subpar single-cell transcriptomes. Initially, we only kept single cells with expressed genes ranging from 500 to 5,000, mitochondrial reads less than 5%, and genes that in more than three cells. Single cell data was normalized by SCTransfrom function in seurat. A principal component analysis was then carried out. After that, data sets from different platform were integrated using the first 20 PCs with Harmony(*1*). Resolution was set to 0.3 to identify clusters. Finally, Doublets were identified by DoubletFinder, and discarded(*2*).

[1] Korsunsky, I. et al. Fast, sensitive and accurate integration of single-cell data with Harmony. Nat. Methods 16, 1289–1296 (2019).

[2] McGinnis CS. et al. DoubletFinder: Doublet Detection in Single-Cell RNA Sequencing Data Using Artificial Nearest Neighbors. Cell Syst. 2019;8(4):329-337.e4.
